# A novel mechanism of microbial attachment: the flagellar pump of *Giardia lamblia*

**DOI:** 10.1101/2024.07.09.602685

**Authors:** Theodore J. Picou, Haibei Luo, Robert J. Polackwich, Beatriz Burrola Gabilondo, Ryan G. McAllister, David A. Gagnon, Thomas R. Powers, Heidi G. Elmendorf, Jeffrey S. Urbach

## Abstract

The ability of microbes to attach to biological and inert substrates is a necessary prerequisite for colonization of new habitats. In contrast to well-characterized mechanisms that rely on specific or non-specific chemical interactions between microbe and substrate, we describe here an effective hydrodynamic mechanism of attachment that relies on fluid flow generated by the microbe. The microbe *Giardia lamblia*, a flagellated protozoan parasite, naturally attaches to the microvilliated surface of the small intestine but is also capable of attaching indiscriminately to a wide range of natural and artificial substrates. By tracking fluorescent quantum dots, we demonstrate a persistent flow between the parasite and substrate generated by a pair of *Giardia* flagella. Using both experimental measures and computational modeling, we show that the negative pressure generated by this fluid flow is sufficient to generate the previously measured force of attachment. We further show that this dynamically-generated negative pressure allows *Giardia* to attach to both solid and porous surfaces, thereby meeting the real-world demands of attachment to the microvilliated surface of intestinal cells. These findings provide experimental support for a hydrodynamic model of attachment that may be shared by other ciliated and flagellated microbes.

## Introduction

Many microbes adhere to surfaces as a first step in colonization of a new niche. Robust attachment is particularly important because many environments experience fluid flow that removes unattached microbes. For example, pathogens are subjected to physiological fluid movement at mucosal surfaces in animal hosts and free-living microbes are subjected to water flow in aqueous environments. While many microbes are quite specific in their attachment site and/or substrate, others can attach indiscriminately to a wide variety of surfaces. Yet in the instances studied to date, the mechanism of attachment is primarily chemical adhesion. For many microbial pathogens, this mechanism is a specific ligand-receptor interaction between molecules on the surface of the microbe and the host cell, while for other pathogens and freeliving microbes, the chemical adhesion is a more general charge-based attraction. This raises the question of whether novel mechanisms of attachment exist that are not reliant primarily on adhesion forces.

*Giardia lamblia*, also known as *Giardia intestinalis* or *Giardia duodenalis*, is a member of the Diplomonadida order. Many Diplomonadida are pathogens of fish, reptiles, birds, and mammals. *Giardia* has two life cycle stages: a cyst stage which is ingested by its host and a metabolically active trophozoite stage which colonizes the intestinal tract. During infection, the trophozoite stage colonizes the surface of the small intestine and attaches to its wall, which is an uneven and pliable surface of intestinal epithelial cells that are covered with microvilli (1, 2). The trophozoite, sketched in Fig. 1a, is 12−15*µ*m long, 5−9*µ*m wide, and ≈ 2*µ*m high. These heterotrophic protists share several notable morphologic features including two nuclei, eight flagella, and a highly structured ventral surface. The four pairs of flagella (anterior, ventral, posteriorlateral, and caudal) play well-defined roles in swimming and steering (3, 4).

**Fig. 1.**
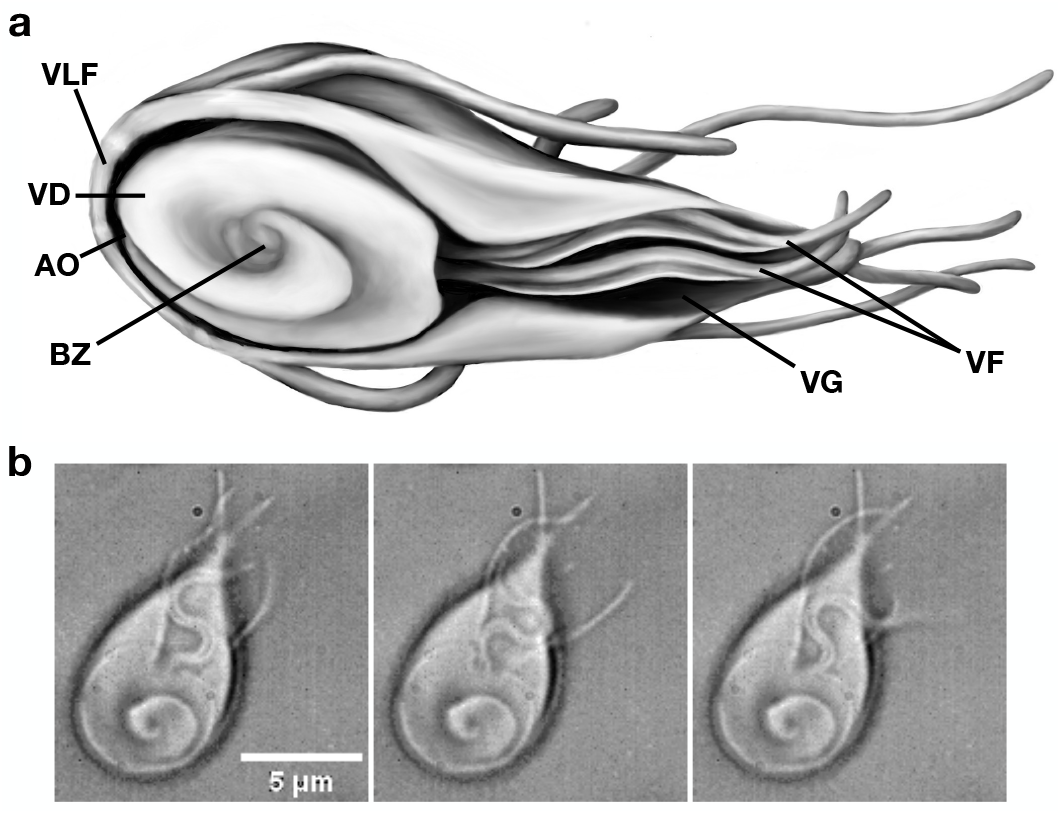
Morphology of attached *Giardia*. (a) Sketch of *Giardia*, indicating the position of the ventrolateral flange (VLF), ventral disk (VD), anterior opening (AO), bare zone (BZ), ventral groove (VG), and ventral flagella (VF). (b) Three frames from a DIC movie (Supplementary Movie SM1) of *Giardia* attached to a glass slide, showing the VF pair beating in the space defined by the VG (60x, 50 ms between frames).

The ventral surface of the trophozoite is dominated at the anterior end by a large domed structure, termed the ventral disk (VD in Fig. 1a), formed by a spiral array of microtubules with a circular opening at their center, the bare zone (BZ). The spiral nature of the array leaves a small gap, the anterior opening (AO) (5–7). In the tapered posterior half of the cell, the ventral surface is concave, forming a chamber with the attachment surface. This structure, termed the ventral groove (VG), houses the ventral pair of flagella (VF) that beat continuously in a modified sinusoidal waveform (Fig. 1b, Supplementary Movie SM1, reviewed in (8)).

In contrast to the multifaceted and dynamic environment *Giardia* experience in the gut, *Giardia* in the laboratory grow attached to glass or plastic surfaces in stationary anaerobic cultures, and studies of *Giardia* attachment have traditionally used these hard, smooth substrates (9, 10). Using an assay to determine the buoyant force necessary to cause cells to detach, *Giardia* were shown to attach equally well to substrates that are chemically diverse with no significant difference in attachment force between uncoated glass and glass coated with poly-L-lysine, a teflon-like coating, or polyethylene glycol (PEG) coating (9). Thus the receptor-ligand binding or other biochemical mechanisms that normally mediate attachment in biological contexts are not required for fully effective *Giardia* attachment.

Notably, while attached, the ventral flagella beat continuously and steadily in the chamber defined by the ventral groove with frequencies ranging from 2.5 to 18 Hz (11–13) (see also Supplementary Figure S2b). Figure 1b shows representative snapshots of a cell attached to a microscope cov-erslip (see also Supplementary Movie SM1). The spiral microtubule array that forms the ventral disk is prominent (although the dome-like morphology is not evident in the twodimensional image). The two flagella are confined by the ventral groove throughout the beating period, and while not evident in the optical images, electron microscopy images show that the flagella are tightly confined on the dorsal side by the ventral groove (8, 14, 15). Aided by the fin-like projection on the ventral side (Fig. 1, (8)), this confinement will ensure that the fluid in each half-wavelength is effectively trapped, and therefore largely removed from the groove with each cycle, creating something approaching a positive-displacement microscale pump. Another notable feature of *Giardia* attached to hard surfaces is the protrusion of cytoplasm and cell membrane at the bare zone (BZ), the hole at the center of the ventral disk, into the space below the disk, coming into contact with surface of attachment (Supplementary Movie SM2, also (10, 15)).

Early efforts to resolve the conundrum of *Giardia* attachment based on these observations came from a series of papers published by David Holberton in the 1970s (11, 16, 17). Holberton proposed a hydrodynamic model of attachment in which these flagella generate fluid flow between the ventral surface of the cell and substrate (the microvilli of the intestinal epithelial cells or the glass/plastic of a tissue culture container), and that this flow in turn generates a pressure differential that causes the cell to attach to the substrate. Unfortunately, Holberton lacked the experimental methodology to test his hypothesis. The hydrodynamic model was slowly abandoned over the subsequent 40 years, but no serious alternative has emerged to take its place that can explain the full range of observed attachment behaviors.

More recent work has shown that the dynamic morphology of the ventral disk (14, 18) and of the ventrolateral flange (15) as *Giardia* interacts with a substrate play important roles in attachment, but, as discussed below, passive mechanisms controlled by the disc morphology alone are insufficient to maintain the sustained pressure differential necessary to maintain attachment. Indeed, a recent study using single-cell force spectroscopy provides support for an active suction mechanism (19).

In this paper, we extend this work by presenting multiple lines of complementary evidence that provide the first direct quantitative support for a hydrodynamic model of attachment. Specifically, we show that the beating of the ventral flagella of attached *Giardia* generates a continuous flow underneath the attached cells, that the pressure differential generated by the flow is sufficient to account for the observed attachment of *Giardia* to surfaces with a wide range of chemistries and topographies, and finally we show that quantitative measurements of the negative correlation between substrate permeability and attachment strength are consistent with the proposed attachment mechanism.

### Demonstration of fluid flow at the attachment interface

To study fluid flow beneath the ventral surface of *Giardia*, we performed particle tracking velocimetry (PTV) on cells attached to glass surfaces using fluorescent particles imaged with both total internal reflection fluorescence microscopy (TIRF), which provides high-resolution imaging at the substrate surface, and spinning disk confocal microscopy, which provides high-speed image capture. Unattached cells swim continuously, making attached cells, which are motionless except for the beating flagella, easy to identify. Beads larger than 20 nm observed in the vicinity of attached cells were consistently excluded from the ventral side the cell. With 20 nm fluorescent beads and 10-20 nm fluorescent quantum dots, however, we observed that particles would occasionally enter the area under the ventral disk. Many of these particles tracked along a very specific pathway, entering at a precise point just off center at the anterior end of the disk, traveling in an arc under the disk, emerging out from under the posterior end of the disk, and traversing the ventral groove (Fig. 2a, Supplementary Movie SM3). This directed motion contrasts with particles just beyond the cell periphery, which show directionless Brownian motion, indicating no significant fluid flow (Supplementary Fig. S1). Particle displacements determined from particle tracking of several cells are shown in Fig. 2b. The particle motions reveal a consistent flow towards the posterior of the cell, with some variability in the point of entry into the region under the disk, most likely reflecting cells that were momentarily not tightly adhered to the surface of the glass coverslips. The range of particle speeds measured in different regions of the cell is displayed in Fig. 2c. with a fluid velocity under the disk that is relatively steady and in the range of 5–15 *µ*m/s, and highly variable in the range of 10–25 *µ*m/s in the ventral groove.

**Fig. 2.**
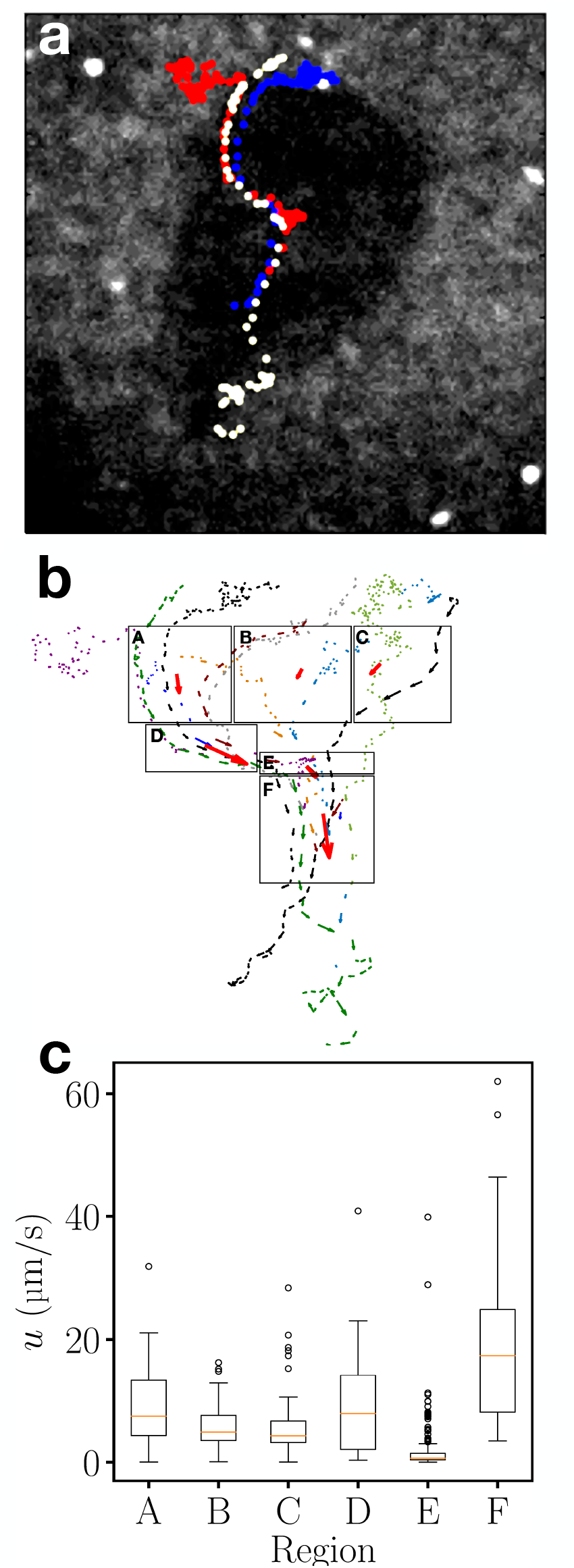
Trajectories of tracer particles. (a) Tracks of 3 fluorescent quantum dots (shown in red, blue, and yellow) under the surface of a cell. (Individual trajectories shown in Fig. S1)). The teardrop shadow of the cell is faintly visible (60x, 20 fps). (b) Overly of particle tracks from multiple cells. All tracks were rotated and scaled to an average-sized trophozoite aligned vertically. (c) Particle speeds from the boxed regions in (b). Boxes A, B and C cover the anterior half of the parasite and contain quantum dots entering under the ventral disk. Boxes D and E record the path of quantum dots exiting the posterior of the ventral disk. Box F shows the path of quantum dots in the ventral groove.

Previous research has reported that 90–200 nm ZnO beads are limited to the periphery of attached cells (13), consistent with our observation that particles with diameters > 20 nm do not pass underneath the ventral disk. This is also consistent with high resolution TIRF mapping of the ventral surface of the cell, showing that ventral disk makes a closed circle in the TIRF images (Supplementary Movie SM2, (15)). However, TIRF illuminates a region up to ∼100nm above the glass surface (20, 21), allowing for the possibility of sufficient space between the cell and substrate for fluid flow and transport of very small particles. Interestingly, some of the particle trajectories show pauses at the boundary between the ventral disk and the ventral groove (e.g Fig. 2a, red particle), consistent with a thin flowing layer close to the glass surface passing under the disk and nearly stationary fluid trapped by the disk above it.

### Modeling

The experimentally-observed fluid flow suggests a ‘flagellar pump’ model for *Giardia* attachment to smooth surfaces. The beating of the flagella in the ventral groove continually draws fluid from underneath the ventral disk into the ventral groove. This requires a continual flux of fluid beneath the disk, which we model as a flow through the small opening on the anterior end at the junction of the microtubule spiral that makes up the ventral disk (AO in Fig. 1) when *Giardia* is attached to a solid surface. Any small gaps between the ventral surface of the cell and the substrate would produce qualitatively similar behavior and are not explicitly included in the model. The small opening represents a high impedance for fluid flow and therefore will generate a large, highly localized pressure gradient. As a result, the pressure in the ventral disk will be significantly below ambient pressure. In contrast to the small opening, the region under the ventral disk itself provides a relatively large space for fluid flow, so there will be an approximately uniform low pressure present throughout the area under the disk. Thus, we propose that the primary purpose of the disk is not to generate low pressure by itself, but to distribute the flow-generated low pressure over a large area.

We have quantitatively tested this model by numerically solving for fluid flow in a two-dimensional model of *Giardia* with beating flagella. Due to the small length scales of the cell, the flow is dominated by viscous forces in a regime commonly referred to as Stokes or low Reynolds number flow, where the fluid flow is simply a balance between pressure and viscous forces. The model geometry, shown in Fig. 3a and described in Methods, is derived from our own experimental observations of attached cells and numerous images in the literature, including the main features shown in Fig. 1, with a circular barrier in the center of the ventral disk to account for the blockage created by the cytoplasm and cell membrane protruding at the bare zone, as described above. The fluid flow and pressure changes generated by the flagellar motion are calculated by solving the viscous flow equations for low Reynolds number flow with no-slip boundary conditions on all surfaces using a commercial finite element package (Comsol Multiphysics). The amplitude of the flagella waveform grows approximately linearly along the length (13), so that the ratio of the amplitude to the channel width, *h*, is constant. A snapshot of the computed flow and pressure fields is shown in Fig. 3a, for an anterior opening (AO) of width 2*a* = 125 nm and a flagellar wave height of *h* = 0.9 (the largest value of *h* for which the simulation reliably converges).

**Fig. 3.**
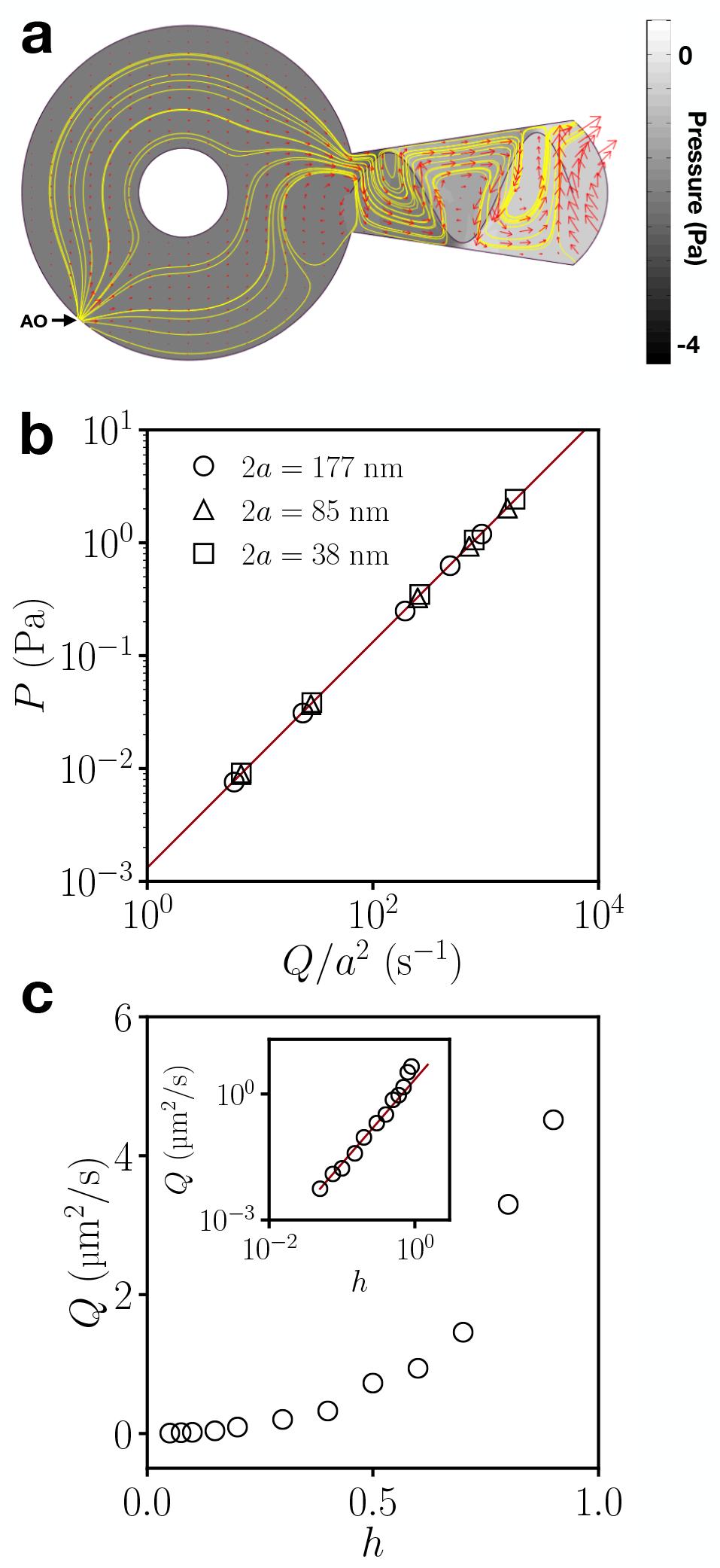
Computational model of fluid-mediated attachment. (a) Snapshot from the 2D computational model. The morphologically accurate geometry includes a circular ventral disk with a small anterior opening (AO) at the bottom left, corresponding to the gap in the region of overlap of the spiral array of microtubules in the ventral disk, and a circular obstruction in the middle, representing the cell protrusion from the bare zone observed in attached parasites. The ventral groove (right) contains ventral flagella beating sinusoidally, pictured for *h* = 0.9 (see Methods) and AO width 2*a* = 125 nm, showing streamlines (yellow), velocities (red vectors), and pressure relative to ambient (greyscale). (b) Pressure versus flux from the computational model for different AO widths (indicated in legend) and wave amplitudes *h* = 0.1, 0.2, 0.5, 0.7, and 0.9. The solid line is a fit to Δ*P* = *CηQ/a*^2^, with C=1.32. (c) Cycle-averaged flux versus wave amplitude. Inset: logarithmic plot of the same data, showing quadratic dependence for small *h*, and faster increase as *h* approaches unity.

The instantaneous fluid velocity (red arrows) and streamlines (yellow lines) are consistent with the experimental observations (Fig. 2). As expected, the low pressure generated at the base of the flagellar pump is spread over the entire area of the ventral disk. The anterior pressure gradient is confined to a very small region around the anterior opening. The general features of the flow field do not change with the phase of the flagellar wave, but fluid flux through the cell and the magnitude of the pressure drop do show modest variation over the course of the cycle (all flux and pressure values reported here averaged over one full period).

The two primary components of this model for attachment, the behavior of the flagellar pump and the pressure drop across the anterior opening of the ventral disk, can be understood directly from the equations for Stokes (low Reynolds number) flow. For a given 2D flux *Q*, the pressure drop generated by the flow through a small opening of half-width *a* is Δ*P* = *CηQ*/*a*^2^, where C is a constant of order unity (equal to 8/*π* for the ideal case of an opening in an infinite wall (22)). Fig. 3b summarizes the results of model calculations for a range of values of anterior opening half-width *a* and flagellar wave height *h*. The expected linear relationship between *P* and *Q*/*a*^2^ is satisfied in all cases, indicating that the pressure under the disk can be understood as the pressure differential generated when the flagellar pump draws fluid through the small anterior inlet (or other gaps in the seal between *Giardia* and substrate).

The flagellar pump is comprised of a pair of flagella beating in a highly confined space. The effects of flagellar waves have been extensively analyzed in the context of cell motility, both in the absence of nearby boundaries and swimming close to walls.(23–26). We have extended that analysis to a simplified model of beating flagella in a confining channel with a small inlet (see Supplementary Material). The linearity of the Stokes equations implies that the flux is the sum of the flagellar-driven flow and the back-flow arising from the pressure gradient. For small wave amplitudes and small inlet openings, we find that the pressure at the base of the beating flagellum, and therefore the pressure under the ventral disk and the resulting attachment force, is proportional to the square of the wave amplitude, consistent with the results of the simulation (Fig. 3c). At larger amplitudes, however, the flux grows more quickly with wave amplitude (inset, Fig. 3c), although even at *h* = 0.9 is much less than expected for a positive displacement pump (≈ 20*µ*m^2^/s for the parameters used in Fig. 3c).

The attachment force predicted by the model can be determined by integrating the pressure drop over the area of the cell, with the largest contribution coming from the ventral disk, and thus is on the order of Δ*P* × *A*_*disk*_, where 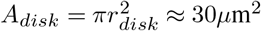 is the area covered by the ventral disk. Δ*P* is very sensitive to the parameters of the model (and in particular to the size of the anterior opening), but is of the order of a few Pa for *h* = 0.9, and would therefore be a few tens of Pa for *h* = 1, resulting in an attachment force Δ*P* × *A*_*disk*_ ∼ 1 nN, slightly less but stil comparable to the attachment forces of ∼ 2*nN* measured by centrifugation (1) and ∼ 8*nN* measured by single-cell force spectroscopy (19).

### Attachment to Porous Surfaces

In the intestine, *Giardia* attach to a microvilliated epithelial layer, so that the mechanism of attachment must be robust to a ‘leaky seal’ between the cell and the substrate. The hydrodynamic model described above can be used to make specific predictions about how leakage will compromise attachment. To test these predictions, we have developed a method for measuring the ability of attached cells to withstand shear forces and investigated the effect of changing substrate permeability on attachment. We use a rheometer coupled to a high-resolution microscope that enables the visualization of fluorescently-labeled attached cells under precisely controlled shear flow. Using custom image analysis routines, we count the fraction of cells that remain attached as the shear rate is slowly increased. Figure 4 shows results for substrates coated with 50 *µ*m thick layers of polyacrylamide (PA) with different acrylamide concentrations. The ability of the cells to withstand shear flow at the highest PA concentration (7%, blue curve) is similar to that observed on glass (Supplementary Fig. S2a), and roughly consistent with the force of attachment measured by centrifugation (9). The shear stress *τ* experienced by the cells in a shear flow 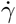 is of order 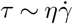. The cell culture media used for these experiments has a viscosity similar to water, *η* ≈ 0.7 mPa·s at 37°C, indicating that cells on substrates with low permeability detach when the shear stress is a few Pa.

**Fig. 4.**
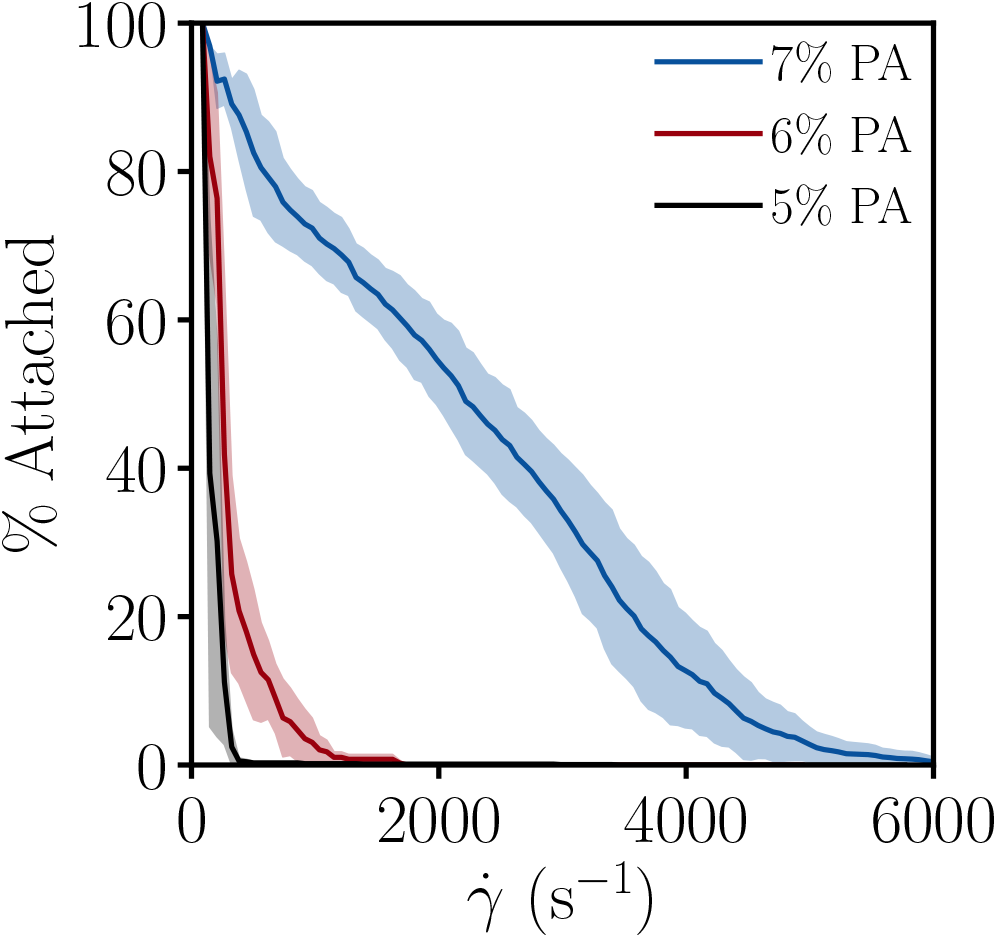
Attachment for substrates of different permeability. The percentage of initially attached cells remaining in place as the shear flow 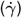 is slowly increased, for substrates coated with thin layers of polyacrylamide (PA) of different concentrations. The solid lines are the average, with the shaded region showing the range for each shear rate (N=3).

Our results, other measurements of flow-induced detachment (27), and the centrifugation (1) and single-cell force spectroscopy (19) experiments show a wide range of attachment forces. This may be due to the almost 10-fold range in ventral flagella frequency that we and others have observed (2-20 Hz; mean ≈ 10 Hz, Supplementary Fig. S2b, also (11–13)), which is linearly related to attachment force in the hydrodynamic model. (Note that the flow assay measures the stress at which cells begin to slide, while the centrifugation and single-cell force spectroscopy assays measured the buoyant force at which cells separate from the substrates. We observe that sliding occurs at shear forces substantially smaller than the detachment buoyant forces.)

We have found that the attachment force is sensitive to the permeability of the substrate. Changing the concentration of acrylamide in the PA layer from 7% to 6% while keeping the ratio of cross-linker constant increases the permeability by about a factor of 2 (from ≈ 3 to 5 ×10^−18^m^2^ (28)), and substantially impairs attachment (compare blue and red curves in Figure 4). Reducing the acrylamide concentration to 5% (black curve) further increases the porosity of the substrate and concomitantly reduces the ability of the cells to remain attached in the presence of shear forces. This behavior is consistent with what would be expected for the proposed hydrodynamic model of attachment: for a given flagellar-driven fluid flux, flow through the ‘leaky’ substrate and into the space beneath the ventral disk will reduce the flow through the small anterior opening and therefore reduce the pressure drop.

The relationship between the substrate permeability *κ*, pressure variations, and the flow through the substrate can be estimated using Darcy’s Law,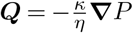. From dimensional considerations, we expect that the total flux through the substrate will be given by *Q*_substrate_ = *CκL*Δ*P*/*η*, where *L* is the perimeter of the cell, defining the region of reduced pressure. We derived an approximate solution with the relevant boundary conditions and found that *C* ≈ 2 (see Supplementary Materials). For the 6% PA substrate, we would therefore predict a flow through the substrate of *Q* ≈ 2 × 10^−19^*m*^3^/Pa. The flagellar driven flux in our 2D model discussed above is of order 10^−11^m^2^. As discussed below, there are a number of issues that complicate extending the model to three dimensions, but an order of magnitude estimate of the flux can be obtained by multiplying the 2D flux by the height of the ventral groove, ≈ 100 nm, so *Q*_3*D*_ ∼ 10^−18^m^3^, which is of the order of the leakage flow predicted by this 10 Pa pressure differential.

The quantitative agreement between the predicted and measured permeability necessary to compromise attachment provides strong support for the hydrodynamic model of attachment. The details, however, suggest that the situation is more complicated than the specific model outlined above. In particular, the data in Fig. 4 show that a factor of three increase in permeability generates an order of magnitude reduction in the shear stress necessary to detach cells. The shear flow assay, however, assesses sliding, rather than directly measuring the force due to the pressure differential, and it is quite possible that the lateral traction force available to resist sliding is not simply proportional to the force normal to the surface produced by a pressure differential. In addition, reduction in the pressure differential could produce additional paths for fluid flow (e.g. by enabling the formation of gaps between the ventral surface and the substrate), producing positive feedback as the permeability compromises the pressure differential. The behavior of the flagellar pump itself will also be affected by substrate permeability, as the shear rates and pressure gradients in the vicinity of the beating flagella are large. Finally, we note that the frequency of the flagella does also change modestly but systematically with substrate porosity, increasing by about 40% on the highest porosity substrates relative to glass (Supplementary Figure S2c).

## Discussion

The mechanism by which *Giardia* attaches to the host intestine has recently garnered interest after almost 50 years of relatively little attention (10, 14, 15, 18, 27, 29). The results we describe in this paper clearly demonstrate that *(a)* the beating of the ventral flagella of attached *Giardia* generate flow underneath the attached cells, *(b)* the pressure differential generated by the flow can account for the observed attachment of *Giardia* to surfaces with a wide range of chemistries and topographies, and *(c)* quantitative measurements of the impact of substrate permeability on attachment are consistent with the proposed attachment mechanism and are sufficient to allow *Giardia* to attach to the semi-porous surface of microvilliated intestinal cells. Thus, this work provides a definitive answer to the long-standing question of the active mechanism primarily responsible for maintaining suction underneath the *Giardia’s* ventral disk. Given the complexity of the natural environment in which *Giardia* attaches in the small intestine, the parasites most likely employ a combination of suction, clutching, and electrostatic attachment forces in vivo, as suggested by others (e.g., (27)).

While the structural similarity of the disk to a macroscopic suction cup might suggest a passive suction mechanism, cell membranes are permeable to water, and water is nearly incompressible. As a consequence, even in conditions where the ventral disk could form a tight seal with the substrate unlikely against the intestinal wall diffusive water transport would compromise attachment nearly instantaneously in the absence of an active mechanism for continuously removing water from under the ventral disk (details in Supplementary Material). Intriguingly, the protrusion of the cell membrane through the bare zone observed by us and numerous other groups e.g., (10,14,15,18,29) only in well-attached parasites is most easily explained by a low-pressure zone beneath the ventral disk. Indeed, structural integrity of the ventral disk itself is essential for attachment (14, 18). Our experimental discovery of fluid flow between the ventral disk of attached parasites and the substrate is explained by the computational model that identifies the waveform of the ventral pair of flagella in the ventral groove as the pump responsible for the fluid flow and consequent pressure differential that drives attachment. Our finding that *Giardia* attaches less well to increasingly porous polyacrylamide gels demonstrates what happens when the influx of fluid exceeds the capacity of the flagellar pump, and we note that a different group observed similar results for *Giardia* attachment under flow on micropatterned surfaces (27).

Similarly, it follows that disrupting either the ventral groove morphology or the ventral flagella waveform should decrease the unidirectional fluid displacement and compromise the strength of attachment. This prediction is supported by the work of Hardin and colleagues who found that disrupting the structural integrity of the ventral groove by knockdown of a prominent protein, Flangin, diminished the force of attachment, rendering parasites still able to attach in stationary culture but less able to attach in a flow cell (15). Earlier, House and colleagues had used morpholinos against two different proteins, PF16 and alpha2-annexin, associated with the ventral flagella, to disrupt the regular and rapid sinusoidal waveform (10). The effects on attachment were evident but less dramatic, leading them to conclude that the ventral flagella were unimportant in attachment. However, we note that a close analysis of the videos they provide show that while the waveform is strikingly altered, the ventral flagella movements still fulfill the conditions set in our computational model i.e., the flagella move to sweep the ventral groove fully in a manner that would expel fluid and we therefore see their data as again supportive of the model we present in this paper

One limitation of the 2D model we describe is that it does not capture the fact that the ventral flagella of *Giardia* are sandwiched between the ventral surface of the cell and the substrate to which the cell has attached (16). The existence of gaps between the flagella filament and those surfaces would mean that not all of the fluid in the ventral groove will be removed with each cycle (so that the flagellar pump is not truly ‘positive displacement’). In addition, the motion of the flagella in close proximity to those surfaces will generate large shear rates and therefore large amounts of viscous dissipation. As mentioned above and shown in Fig. 1a, the ventral flagella possess unique structures on their ventral side, called fins (16). These fins likely serve to minimize the gaps, acting like windshield wipers to push fluid that is close to the surface of attachment, so that leakage would be minimal. A somewhat analogous approach is employed by leucon sponges, filter-feeders that employ pumps in choanocyte cells based on flagella with a ribbon-like morphology (30, 31).

Another limitation of the model is that the flagellar waveform and the configuration of the parasite are taken as fixed, while in reality the large forces generated by forcing fluid flow in a tightly confined environment will constrain the flagellar beating and deform cell surfaces. For example, the portion of the ventral disk that separates the low pressure region under the disk from the ventral groove will experience localized forces that will vary with the phase of the flagellar beating, and could cause deformations that would modulate the impedance between the region under the disk and the groove. Finally, for simplicity we consider only the impedance arising from flow through the anterior opening (Fig. 3), while in reality the ventrolateral flange (Fig. 1, VLF) can be in close proximity to the attachment surface and therefore can produce additional, likely variable impedance. Nonetheless, the simplified model seems to capture the primary features observed when *Giardia* attach to rigid surfaces.

More generally, while this study has focused on understanding *Giardia* attachment in culture, the mechanism we describe will also be effective in the intestine that presents an uneven and pliable surface of viscoelastic mucus layers and microvilli (1, 32). Thus the insights gained from this work have potential relevance to a wide range of biophysical systems, including other flagellated protists (33), other situations where filaments beat near boundaries, and biomimetic applications for active pumping and attachment to diverse surfaces.

## Methods

### Cell culture and labeling

*Giardia lamblia* trophozoites (isolate WB and in separate experiments, isolate GS) were maintained anaerobically in borosilicate glass tubes or polystyrene flasks. Parasites were grown at 37°C in modified TYI-S-33 media (Keister, 1983), where 0.024 M sodium bicarbonate replaced the traditional phosphate buffer solution. For all experiments, *Giardia* cultures were grown to mid-log phase (∼ 80% confluency).

For labeling of nuclei with Syto-16, cells first were collected by chilling and centrifugation and resuspended in sterile PBS supplemented with 6 *µ*M Syto16. Cells were labeled for 30 minutes at 4°C with constant rotation in the dark, collected again by centrifugation, and resuspended in fresh cold PBS. For cell surface labeling, culture media was decanted and fresh media supplemented with 10 *µ*g/ml wheat germ agglutinin conjugated to Alexa-488 (Invitrogen) was added for a 2-hour incubation at 37°C. Cells were washed three times in culture media, chilled to detach, collected by centrifugation, and resuspended in a small volume of culture media.

### TIRF microscopy

Prior to TIRF imaging, the uniformity of cell surface labeling was determined by confocal microscopy. Labeled *Giardia* were resuspended in cold media, added to the glass coverslip of a Lab-Tek chamber slide, and placed in the 37°C incubation chamber of an Olympus IX81 Total Internal Reflection Microscope. As parasites began to attach to the glass coverslip, focus level and incident angel of the argon 488nm laser were adjusted to obtain the optimal TIRF images. Images were taken at 60X using a Plan Apo 60X oil/N.A. 1.45, WD 0.1 mm with correction collar objective, captured using a Hamamatsu C9100-12 EM 512Å 512 back-thinned CCD digital camera driven by IP Lab Suite software with motion control. Wide-field images corresponding to TIRF images were taken as controls to show the full view of cells. TIRF and wide-field images were analyzed using Metamorph software to quantify fluorescence intensity, an indirect measure of distance between cell and substrate assuming uniform cell surface labeling.

### Particle tracking velocimetry

Cells were resuspended in a buffer of 1X PBS, 10% Ficoll, 4% bovine serum albumin, and a 1:25 dilution of Qtracker 655 non-targeted quantum dots (Invitrogen) for imaging. The Ficoll was added to increase the viscosity (4.8 cP) of the media to minimize Brownian motion, and bovine serum albumin was introduced to prevent the quantum dots from adhering to the glass microscope coverslip. Video was captured at 60X magnification using a Nikon Eclipse TE2000-U spinning disk confocal microscope with an Andor iXon DV887 camera and a temporal resolution of 50 ms using SlideBook (Intelligent Imaging Innovations) microscope imaging software. Particles were tracked with custom Matlab code adapted from (34).

### Finite element computational model

Low Reynolds number fluid flow was modeled using the Comsol Multiphysics modeling and simulation software solving for a deformed moving mesh representing a single continuous line boundary formed by the ventral flagella pair, consistent with the observation that the pair normally beat in unison (Supplementary Movie SM1). No-slip conditions were specified on all of the boundaries except the inlet and outlet, which were defined to be neutral. Instantaneous flow and pressure fields were calculated at 12 time points during a single period of the beating flagella. Following ref. (13), the flagellar boundary is modeled as a traveling wave with a linearly increasing amplitude, *y*(*x, t*) = *h* * *s*(*x*) * sin(*kx − ωt*), where *x* is the linear distance along the centerline of the parasite starting at the anterior opening, *y* is the perpendicular displacement of the flagellum, *ω* is the angular frequency of the flagellar wave, *ω* = 23.6 rad/s, and *k* = 2*π/λ*, with *λ* = 2.66*µm*, and *s*(*x*) = 0.15*x* + 0.7 is the line matching the edge of the ventral groove, so that when *h* = 1 the peaks of the flagellar wave just make contact with the walls of the ventral groove at every point in the beating cycle.

### Polyacrylamide substrates and attachment assay

Glass coverslips were cleaned by washing thoroughly twice, in sequence, ethanol and water. The coverslips were then passed over the inner blue flame of a Bunsen burner (with the side to be activated facing down) to remove any particles and ready the coverslips for activation. A small amount of 0.1 N NaOH was pipetted on the activated sides, spread evenly using a Kimwipe, and air-dried for approximately 10 minutes. 50 *µ*L of Aminopropyl-trimethoxysilane (APTMS, Sigma-Aldrich) was pipetted on the center of the coverslips and spread evenly over their surfaces using the pipette tip. After waiting 5 minutes to react, the coverslips were placed in ddH_2_O for approximately 10 minutes to allow complete removal of the excess APTMS. Following the ddH_2_O wash, coverslips were air dried before being completely covered with 0.5% glutaraldehyde and allowed to sit for 30 minutes. Excess glutaraldehyde was removed, and the coverslips were rinsed in ddH_2_O for 10 minutes before allowing to air-dry.

Solutions of 5, 6 and 7% acrylamide:0.03% bis-acrylamide (Sigma-Aldrich) were allowed to polymerize between the activated surface of hydrophilic coverslips and a hydrophobic glass slide for 25 minutes, using the rheometer to produce a spacing of precisely 50 *µm*. After polymerization, the rheometer tool was raised, the glass slide removed, and the tool lowered again to produce a gap between the tool and the acrylamide layer of 100 *µ*m.

*Giardia* labeled with Syto-16 were added to either uncoated glass slides or to slides with the acrylamide layer and incubated for 10 minutes at 37°C to allow for cell attachment, at which point the rheometer was used to produce a 120 second linear shear ramp starting at 100 s^−1^ and ending at 8000 s^−1^. Images were captured every second, with a 50 ms exposure time, starting from 10 seconds before the beginning of the shear ramp.

### Cell counting

Candidate cells were located in images subjected to band-pass filtering, with centroid location, brightness, and eccentricity used to refine the count. Cells that appeared in the field of view during the measurement were excluded.

## Acknowledgments

We thank Aliza Apple and Jesse Cohen for early research contributions to this project, WeiWei Gerl for collecting flagellar frequency data, and Peris Lopez for creating the *Giardia* sketch (Fig 1). We thank Susette Mueller and Peter Johnson of the Microscopy & Imaging Shared Resource at Georgetown University Medical Center for helpful discussion and technical guidance on the TIRF microscopy, and Peter Olmsted for helpful discussions. T.J.P. was supported by the ARCS Foundation. T.R.P.’s work was supported in part by the National Science Foundation under Grant No. PHY-1066293 and the hospitality of the Aspen Center for Physics. J. S. U. is supported in part by the Georgetown Interdisciplinary Chair in Science Fund.

All data are included in the manuscript and/or supporting information. The code used in this manuscript is stored in GitHub (Link to be added before publication).

