## Supplementary Material for "A novel mechanism of microbial attachment: the flagellar pump of *Giardia lamblia*"

### Supplemental Material, Picou, et al.

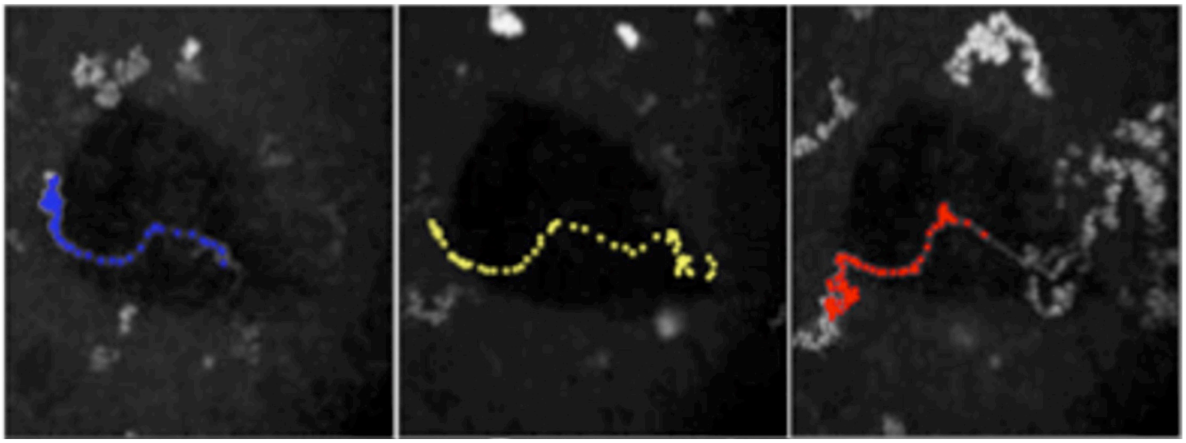

**Fig. S1. Image overlays from Fig. 2a.** Confocal microscopy overlays from the time intervals shown in Fig. 2a. The tracked particles are highlighted, showing directed motion in clear contrast to the random Brownian motion of the quantum dots exterior to the cell.

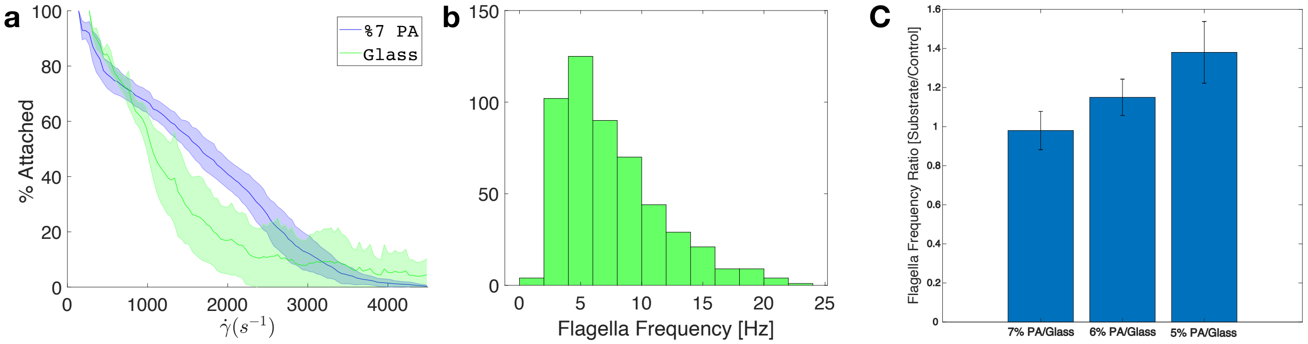


**Fig. S2. Attachment and flagellar beating on glass and polyacrylamide substrates.** (a) The percentage of initially attached cells remaining in place as the shear flow ($\dot{\gamma}$) is slowly increased, on a glass substrate (green) compared with a substrate coated with thin layers of 7% polyacrylamide (PA, blue) The solid lines are the average, with the shaded region showing the range for each shear rate (N=3). (b) Histogram of the average ventral flagellar beat frequency of individual *Giardia* attached to glass coverslips. The average flagellar frequency was determined by observing videos of individual cells over a period of 300 frames ( 3 seconds) and counting the number of cycles of the flagellar wave. (c) The ratio of the mean flagellar beat frequency relative to control (glass) for different concentrations of PA. (Error bars are the standard deviation of the mean frequency ratio cacluclated for each trial, 5 trials of 7%, 3 each for 6 and 5 %, N = 14-49 per trial.)

**Supplementary Movie 1:** High resolution microscopy (DIC) movie of *Giardia* attached to glass coverslip showing periodic beating of ventral flagella (60x, 20 fps). Note images are inverted compared to Fig. 1.

**Supplementary Movie 2:** *Giardia* attached to glass coverslip initially imaged in fluorescence microscopy, where cell outlines are visible, then switching to TIRF microscopy, which only shows cell membrane within a few hundred nanometers of the surface. In the lower left quadrant of the movie, a cell is seen attaching to the surface. Initially, the outline of the cell and the ventral disk is visible, followed by a patch of cell membrane in the center of the disk, indicating cell membrane protruding through the bare zone (see Fig. 1, also Fig. 1 of (1)).

**Supplementary Movie 3:** Spinning Disc Confocal Microscopy movie of fluorescent quantum dots, one of which traverses under the surface of a cell. The teardrop shadow of the cell is faintly visible (60x, 20 fps).

### Supporting Information Text

In Section 1, we consider the impact of diffusive water transport through the cell membrane driven by a pressure difference of the sort hypothesized to be responsible for *Giardia* attachment. In Section 2, we calculate the pressure drop produced by a two-dimensional flagellum beating in a two-dimensional pipe. In Section 3, we consider the effect of a chamber at the base of the flagellum as a model for the Ventral Disk. In Section 4, we estimate the fluid flow through a porous substrate.

#### 1. Cell membrane diffusive transport

In the first section below we consider a passive suction cup and show that, due to the nearly incompressible nature of water, the diffusive flux through the cell membrane, while very small, compromises attachment nearly instantaneously, highlighting the need for some sort of active process to maintain negative pressure beneath attached cells. In the second section, we estimate the diffusive flux that would arise due to the specific pressure differentials hypothsized in the model presented here, and show that they are many orders of magnitude lower than the flux generated by the flagella.

**A. Impact of diffusive transport on a passive suction mechanism.** As an idealized model of a passive suction cup, consider a rigid sealed cup of fixed volume $V_c$ filled with water at standard temperature and pressure. Creating a negative hydrostatic pressure under the cup requires removing a small amount of fluid, thereby reducing the density of the remaining fluid and changing the pressure according to $dP/d\rho = -B/\rho_o$, where $B$ is the Bulk Modulus and $\rho_o$ the density of water at standard temperature and pressure. (Similar considerations would apply to a negative pressure created by slightly increasing the volume of the cup while keeping the quantity of trapped fluid constant.) The bulk modulus of water is $B \sim 2 \times 10^9 Pa$, so the pressure underneath the disk changes dramatically for even very small changes in water content. For example, creating the 10 Pa pressure differential considered above requires removing a volume of only $\Delta V = V_c \Delta P/B$, or about $(5 \times 10^{-7})V_c$. Similarly, the pressure differential will be compromised by a very small influx of water, such as that expected from passive diffusion through the cell membrane.

The volume flow rate across a membrane in response do a hydrostatic or osmotic pressure gradients is given by (2):

$$Q = P_f \{ \frac{S V_w \Delta P}{RT} + \sum_i \sigma_i (\Delta \Pi_i) \} \qquad [1]$$

where $P_f$ is the water permeability coefficient, $S$ is the membrane surface area, $V_w$ is the partial molar volume of water (18 $cm^3/mol$), $\Delta P$ is the hydrostatic pressure differnce across the membrane, $\sigma_i$ is the reflection coefficient of the $i$th solute, and $\Delta\Pi_i$ is the osmalality difference for the $i$th solute, and $R$ the gas constant. The osmotic gradient term usually dominates over the hydrostatic pressure term, but the latter is the term of interest here. We can simplify Eq. 1 by identifying a characteristic pressure $P_o = \frac{RT}{V_w} = \frac{(8J/mol/K)(300K)}{(18\times10^{-6}m^3/mol)} = 1.3 \times 10^8 Pa$, so Eq. 1 becomes

$$Q = P_f \frac{S \Delta P}{P_o}, \qquad [2]$$

keeping only the hydrostatic pressure term. For diffusive transport through lipid bylayers, $P_f \sim 2 - 50 \times 10^{-4} cm/s$ (3), while higher values are usually associated with transport through channels.

To model the effect of pressure-driven flow on a cup made of cell membrane, Eq. 2 can be written in terms of the density variation of the water in the cup, $\Delta\rho = \rho - \rho_o$, using $\Delta P = -B\Delta\rho/\rho_o$. The density of the trapped water, $\rho$, will relax to $\rho_o$ according to

$$d\rho/dt = Q\rho_o/V_c = -\frac{P_f S}{P_o}\frac{B}{V_c}\Delta\rho = -\Delta\rho/\tau \qquad [3]$$

with

$$\tau = \frac{V_c P_o}{(BP_f S)} \qquad [4]$$

Thus the pressure difference will will decay exponentially with time with time constant $\tau$. Taking $P_f = 1 \times 10^{-3} cm/s$, the area of the disk as $S = \pi(2.5 \times 10^{-6} m)^2 \approx 2 \times 10^{-11} m^2$, and the average height as $2 \times 10^{-6}$ m, we find $\tau \sim 10$ ms. While there is considerable uncertainty in the numbers, it is clear that diffusion through the call membrane is sufficient to degrade a the pressure in a static suction cup extremely rapidly.

**B. Estimate of pressure driven transport.** Alternatively, if we presume that there exists some active process that maintains a negative pressure underneath the Ventral Disk, for example by the flagellar driven pump considered in this manuscript, we can estimate the diffusive flux that will be generated by the negative pressure according to Eq. 2. For a pressure difference of 10 Pa = 10 $J/m^3$ and again taking $P_f = 1 \times 10^{-3} cm/s$ and the area of the disk as $S = 2 \times 10^{-11} m^2$, we find

$$Q = (1.5 \times 10^{-24} m^6/J - s)\Delta P \sim 1 \times 10^{-23} m^3/s \qquad [5]$$

as compared to $Q \sim 1 \times 10^{-18} m^3/s$ estimated for the flagellar pump. Thus while the diffusive flux through the membrane is sufficient to compromise a passive suction mechanism, it is orders of magnitude smaller than the flux generated by actively beating flagella.

#### 2. Flagellum in a pipe with pressure gradient

Here we calculate the ratio of flux to pressure drop, or flow conductance, for a two-dimensional flagellum in a two-dimensional pipe. We can think of the flagellum as an infinite sheet with a traveling wave ripple, confined between two parallel plates of infinite extent (Fig. S3). We take the sheet and pipe to be infinite to avoid dealing with end effects. The amplitude of the sheet is small compared to the wavelength, $bq \ll 1$, and the gap $b \ll d$. This simple model for a beating flagellum, without the confining plates, was first considered by Taylor (4). This model has many variants and is discussed in the review (5). The case of swimmer near a gap (with no gradient) is considered by Reynolds (6).

We prescribe the motion of the sheet: material points move up and down in the $y$-direction, and there is no $x$-component of the motion of the sheet. The sheet is held fixed by some internal mechanism that we do not consider—if the sheet were finite in length, we would anchor it at one end. Given the motion of the sheet and the no-slip boundary conditions at the sheet and the walls of the pipe, we must solve the Stokes equations,

$$\eta\nabla^2\mathbf{v} = \nabla p \qquad [6]$$

$$\nabla \cdot \mathbf{v} = 0. \qquad [7]$$

The Stokes equations are appropriate since the Reynolds number is small. For example, if we suppose the characteristic frequency is $\omega \approx 10$ Hz, the characteristic length is $\ell \approx 1\,\mu$m, and the viscosity is that of water, $\eta = 10^{-3}$ Pa-s, then we calculate a Reynolds number of Re $\approx \rho\omega\ell^2/\eta \approx 10^{-5}$, where

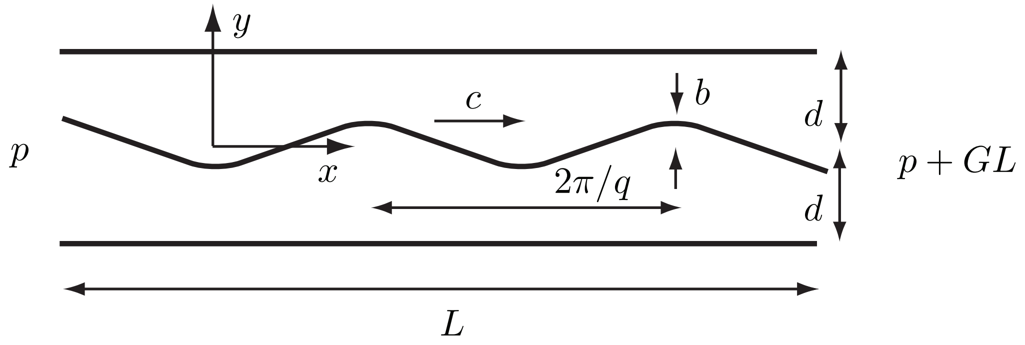


**Fig. S3.** Pump geometry. A traveling wave on a sheet with wavenumber $q$, amplitude $b$, and wavespeed $c$ pumps fluid in the positive $x$-direction down a two-dimensional pipe of diameter $2d$. The sheet is infinite in the $x$ direction and also the direction into the page. The pump works against a pressure gradient $G$.

we used the density of water, $\rho = 10^3\,\text{kg-m}^{-3}$. Note that mucus has a higher viscosity which would make the Reynolds number even lower. Thus, we are justified in taking the Reynolds number to be zero.

Let the subscripts $\pm$ denote the regions above and below the sheet. For example, $\mathbf{v}(x,y,t) = \mathbf{v}_\pm(x,y,t)$ for $y \gtrless 0$. The no-slip boundary conditions are

$$\mathbf{v}_\pm(x,\pm d,t) = \mathbf{0} \qquad [8]$$
$$\mathbf{v}_\pm(x,h(x,t),t) = \dot{h}(x,t). \qquad [9]$$

Note that even though the governing equations are linear in velocity, the no-slip boundary condition at the sheet has a nonlinear dependence on $h(x,t)$. The traveling wave deformation of the sheet is given by $h(x,t) = b\cos(qx - \omega t)$. We will work in units of inverse wavenumber for length and inverse frequency for time, so that $h(x,t) = b\cos(x-t)$. Also, we will expand in powers of $b$ so that at every order we solve a linear problem. Thus, it is convenient to use complex notation and write $h(x,t) = \Re\{b\exp[i(x-t)]\}$. The pumping velocity is second order in $b$ (see (5) and references therein). Since the pressure drop along the flagellum will be set up by the flagellar pumping, we will assume that the pressure gradient $G$ is also second order in $b$.

It is convenient to work in terms of the stream function $\psi$. Eqn. 7 is satisfied by a velocity field which is a curl: $\mathbf{v} = \nabla \times \psi\hat{\mathbf{z}}$, where $\hat{\mathbf{z}}$ points out of the page. The Stokes equation 6 becomes $(\nabla^2)^2\psi = 0$, which we write more simply as $\nabla^4\psi = 0$. The stream function satisfies the biharmonic equation.

**A. First-order problem.** Since the flagellum lies in the middle of the pipe, we will use symmetry to solve the problem. Reflection symmetry implies

$$v_x(x,-y,t) = v_x(x,y,t+\pi) \qquad [10]$$
$$v_y(x,-y,t) = -v_y(x,y,t+\pi). \qquad [11]$$

At first order we have

$$\psi^{(1)}(x,y,t) = \hat{\psi}^{(1)}(y)\mathrm{e}^{i(x-t)}. \qquad [12]$$

Note that linearity and the boundary conditions imply there is no non-oscillatory component. The first order velocity is

$$v_x^{(1)} = \partial_y\psi^{(1)} \qquad [13]$$
$$v_y^{(1)} = -\partial_x\psi^{(1)} = -\mathrm{i}\psi^{(1)}. \qquad [14]$$

Reflection symmetry, described by Eqs. 10–11, implies that $\psi^{(1)}$ is even in $y$:

$$\psi_+^{(1)}(x,y,t) = \psi_-^{(1)}(x,-y,t). \qquad [15]$$

The solution to the biharmonic equation is of the form

$$\psi_\pm^{(1)} = [A\cosh y \pm B\sinh y \pm Cy\cosh y + Dy\sinh y]\,\mathrm{e}^{\mathrm{i}(x-t)}. \qquad [16]$$

There is no uniform component, no shear flow, and no parabolic flow at this order. The reflection symmetry allows us to only consider the part of the problem with $y \geq 0$. The boundary conditions

$$v_+^{(1)}(x,0^+,t) = -\mathrm{i}h(x,t) \qquad [17]$$
$$v_+^{(1)}(x,d,t) = 0 \qquad [18]$$

yield

$$A = b \qquad [19]$$

$$B = \frac{2b(d + \cosh d \sinh d)}{1 + 2d^2 - \cosh 2d} \qquad [20]$$

$$C = -B \qquad [21]$$

$$D = -\frac{2b \sinh^2 d}{1 + 2d^2 - \cosh 2d}. \qquad [22]$$

Since all quantities at first order are proportional to $\exp[\mathrm{i}(x - t)]$, there is no time-average flow, $\langle v_x^{(1)}(x, y, t)\rangle = 0$, where $\langle . \rangle$ denotes the time-average over one period. Thus, the time-averaged flux vanishes to first order. Similarly, the time-averaged force on the flagellum also vanishes to first order in $b$.

The power depends on the product of the velocity and stress at the sheet. Thus, the leading-order power is second order in $b$, and may be deduced from the first-order solution:

$$\langle P_\pm \rangle = \int_0^{2\pi} \mathrm{d}t \, v_{y\pm}(x, 0^\pm, t) f_{y\pm}(x, 0^\pm, t), \qquad [23]$$

where $f_{\pm y} = -p + 2\eta \partial_y v_y$ is the force per unit area on the sheet in the direction of the motion of the material points. Using the $x$-component of the Stokes equation 6 to find $p$, noting that $\langle P_+ \rangle = \langle P_- \rangle$, and using the fact that the average of the square of a cosine over one period is one half, we find that the total power per unit area expended by the sheet is

$$\langle P \rangle = 2 \frac{\eta \omega^2}{q} \frac{b^2 q^2 (2qd + \sinh 2qd)}{\cosh 2qd - 1 - 2q^2 d^2}, \qquad [24]$$

where we have reinstated the dimensions. Although we do not expect the wavelength $2\pi/q$ to be long compared to the diameter $d$, it is convenient for the purpose of illustration to simplify the formulas by working in the 'lubrication' limit $qd \ll 1$, in which

$$\langle P \rangle \sim 12 \frac{\eta \omega^2 b^2}{q^2 d^3}. \qquad [25]$$

The power diverges as $qd \to 0$ since as $q$ gets smaller and smaller, the incompressibility constraint implies that the $x$-component of the velocity gets larger and larger, leading to more and more dissipation.

**B. Second-order problem.** Ultimately we want the time-averaged flux $Q$ down the pipe. Thus we must find $v_x^{(2)}$ using

$$-\partial_x p^{(2)} + \eta \nabla^2 v_x^{(2)}. \qquad [26]$$

The boundary condition at the sheet to second order is found by expanding $\mathbf{v}(x, h, t) = (0, \dot{h})$:

$$v_x^{(2)}(x, 0, t) = -h \partial_y v_x^{(1)}(x, 0, t). \qquad [27]$$

Note that the symmetry of this boundary condition implies that $v_x^{(2)}(x, y, t)$ is even in $y$. If we average 26 over a period (in time or space $x$, it does not matter), we eliminate all $x$-derivatives of oscillatory components of the equation, and we are left with

$$-G + \eta \partial_y^2 \langle v_x^{(2)} \rangle = 0, \qquad [28]$$

with solution $\langle v_x^{(2)} \rangle = E + Fy + Gy^2/(2\eta)$. The boundary conditions 27 and $v_x^2(y = d) = 0$ imply

$$E = \frac{b^2(d^2 + \sinh^2 d)}{\cosh 2d - 2d^2 - 1} \quad [29]$$

$$F = -E/d - Gd/(2\eta). \quad [30]$$

For the flux, note that to second order in $b$,

$$Q_+ = \int_h^d \mathrm{d}y\, v_{x+}(x, y, t) \quad [31]$$

$$\approx \int_0^d \mathrm{d}y\, v_{x+}^{(2)}(x, y, t) - hv_{x+}^{(1)}(x, y = 0^+, t) \quad [32]$$

$$\approx \int_0^d \mathrm{d}y\, v_{x+}^{(2)}. \quad [33]$$

Using $\langle Q \rangle = \langle Q_+ + Q_- \rangle = 2\langle Q_+ \rangle$, and reinstating dimensions, we find that the flow conductance for a flagellum in a pipe with pressure gradient $G$ is

$$\langle Q \rangle = \omega b^2 \frac{qd(q^2d^2 + \sinh^2 qd)}{\cosh 2qd - 1 - 2q^2d^2} - \frac{Gd^3}{6\eta} \quad [34]$$

$$= \omega b^2 F(qd) - \frac{Gd^3}{6\eta}. \quad [35]$$

Equation 35 is the main result of this section. The first term, which is always positive, is the flow generated by the flagellum, and is proportional the frequency and the amplitude squared, with a relatively weak dependence on the ratio of the wavelength to the channel width (through $F(qd)$). The second term is the backflow forced by the pressure gradient. Note again that we have assumed that the pressure gradient is $\mathcal{O}(b^2q^2)$. If we had assumed that the pressure gradient were zeroth order in $bq$, then the pressure-driven flow would dominate the flagellum-driven flow (which would also depend on $G$, since the zeroth-order flow would enter the boundary conditions at first order).

We must be careful if we want to consider the lubrication limit of Eq. 35. The pressure gradient $G$ in our problem is ultimately determined by the flagellar pumping. Thus, if we take $qd \to 0$, we must adjust $G$ so it has the same order as the flagellar-driven flow. With this scaling, the lubrication limit of Eq. 35 is

$$\langle Q \rangle \sim \frac{3\omega b^2}{qd} - \frac{Gd^3}{6\eta}, \quad [36]$$

#### 3. Flagellum attached to a nearly closed cavity

If the flagellar pump is connected to a closed cavity at its base (modeling the Ventral Disk (VD) with no Anterior Opening (AO)), the next flux will be zero, and the two terms on the right hand side of eq. 35 will be equal in magnitude, and the pressure at the base of the flagellum will be reduced from atmospheric pressure by

$$\Delta P = GL = \eta\omega b^2 F(qd) 6L/d^3. \quad [37]$$

This represents the upper limit of the pressure drop in the VD generated by the flagellar pump in the regime where eq. 35 is valid.

We can model the effect of the AO by considering a small opening in the cavity, where the flow conductance through a hole of radius $a$ in two dimensions (a slit of width $2a$ in three dimensions) is

$Q/\Delta P = \pi a^2/(8\eta)$, where $Q$ is the flux and $\Delta P$ is the pressure drop (7). Assuming this flux into the disk is balanced by a net flux out from the pump, eq. 35 becomes

$$Q = \Delta P \pi a^2/(8\eta) = \omega b^2 F(qd) - \frac{\Delta P d^3}{6\eta L} \quad [38]$$

or

$$\Delta P = \frac{\eta\omega b^2 F(qd)}{k_i^{-1} + k_b^{-1}} \quad [39]$$

where $k_i = 8/\pi a^2$ is the inlet impedance, and $k_b = 6L/d^3$ is the backflow impedance. For sufficiently small AO, $k_i >> k_b$, and the pressure in the VD is approximately given by eq. 37, and is therefore independent of the AO, and consistent with the quadratic scaling of $Q$ and $P$ with $h$ observed in the computational model presented in the main text.

#### 4. Leakage through a porous substrate

Consider an infinite half-space $z < 0$ containing a porous material that follows Darcy's law, $\vec{v} = \frac{\kappa}{\eta}\nabla P$ where $\kappa$, is the substrate permeability. The low pressure region represented by the ventral disc is modeled by applying the boundary conditions $p = -p_o$ for $r < a$, and $p = 0$ for $r > a$. Assuming incompressibility, the equation for $v_z$ can be calculated by a Fourier-Bessel expansion:

$$v_z = \frac{p_o \kappa a}{\eta} \int_0^\infty J_1(qa) J_0(qr) e^{qz} d(qa)$$

The flux through the region $r < a$ at $z = 0$ is $Q = 2\pi \int_0^a r v_z dr$, so

$$Q = \frac{2\pi p_o \kappa a}{\eta} \int_0^\infty J_1(u) J_1(u) du$$

or

$$k^{-1} = Q/p_o = \frac{2\pi a \kappa}{\eta} \int_0^\infty J_1(u) J_1(u) du$$

This agrees with the dimensional analysis result presented in the text, with

$$C = \int_0^\infty J_1(u) J_1(u) du$$

The integral diverges, so the upper limit must be cutoff, $u_{max} = (a2\pi/w)$, where $w$ is an estimate of width of the wall at r=a. This makes physical sense, because an infinitely thin wall with a fixed pressure difference would imply an infinite flux at the wall. Using $w$=100 nm and radius $a = 3$ microns, $u_{max} \sim 200$. Using Mathematica to calculate the definite integral we find $C \approx 1.895$. Empirically, C grows as $\log u_{max}$, so the result is relatively insensitive to the chosen cutoff.
